# The diversity and multiplexity of edge communities within and between brain systems

**DOI:** 10.1101/2020.05.05.067777

**Authors:** Youngheun Jo, Farnaz Zamani Esfahlani, Joshua Faskowitz, Evgeny J. Chumin, Olaf Sporns, Richard F. Betzel

## Abstract

The human brain is composed of regions that can be grouped into functionally specialized systems. These systems transiently couple and decouple across time to support complex cognitive processes. Recently, we proposed an edge-centric model of brain networks whose elements can be clustered to reveal communities of connections whose co-fluctuations are correlated across time. It remains unclear, however, how these co-fluctuation patterns relate to traditionally-defined brain systems. Here, we address this question using data from the Midnight Scan Club. We show that edge communities transcend traditional definitions of brain systems, forming a multiplexed network in which all pairs of brain systems are linked to one another by at least two distinct edge communities. Mapping edge communities back to individual brain regions and deriving a novel distance metric to describe the similarity of regions’ “edge community profiles”, we then demonstrate that the within-system similarity of profiles is heterogeneous across systems. Specifically, we find that heteromodal association areas exhibit significantly greater diversity of edge communities than primary sensory systems. Next, we cluster the entire cerebral cortex according to the similarity of regions’ edge community profiles, revealing systematic differences between traditionally-defined systems and the detected clusters. Specifically, we find that regions in heteromodal systems exhibit dissimilar edge community profiles and are more likely to form their own clusters. Finally, we show show that edge communities are highly personalized and can be used to identify individual subjects. Collectively, our work reveals the pervasive overlap of edge communities across the cerebral cortex and characterizes their relationship with the brain’s system level architecture. Our work provides clear pathways for future research using edge-centric brain networks to investigate individual differences in behavior, development, and disease.

## INTRODUCTION

The human brain is a complex network made up of functionally and structurally interacting neural elements [1–3]. Traditionally, brain networks are represented using models in which nodes and edges are defined as regions and the magnitude of their correlated activity, i.e. functional connectivity (FC), respectively [4–6]. This node-centric model emphasizes interactivity among pairs of nodes and has been especially useful in cognitive and network neuroscience, where inter-individual variation has been linked to subjects’ cognitive [7], disease [8], and developmental states [9].

Among the most salient features of node-centric functional networks is their decomposability into subnetworks called “modules” or “communities” [10–13]. In general, networks with modular structure are evolvable [14, 15], are capable of supporting complex dynamics [16], can buffer perturbations, and facilitate cost-effective embedding in three-dimensional space [17]. In the case of human brain networks, the boundaries of modules delineate patterns of task-evoked activity [18] and correspond closely with known cognitive and functional systems [10, 11]. This is true even when modules are estimated under task-free or resting-state conditions. This observation has prompted the hypothesis that modular structure is a key feature for supporting specialized brain function [19].

In virtually every application, the brain’s modular structure is estimated using node-centric FC, which results in a mapping of nodes (brain regions) to modules [20]. Recently, we proposed a novel edge-centric model for representing pairwise functional interactions among a network’s edges [21, 22]. Although node and edge FC (nFC and eFC) are generated from identical fMRI time series, the two constructs provide complementary insight into brain network organization and operation. Whereas nFC measures the extent to which the activity of one brain region fluctuations with the activity of another, eFC unwraps those co-fluctuations across time, first yielding moment-by-moment accounts of the co-fluctuations between pairs of brain regions (edges) and then assessing the similarity between pairs of cofluctuation time series [22].

Similarly, compared to the modular structure of nFC, the modules estimated from eFC provide complementary information about the brain’s system-level organization. Clustering nFC results in a partition of nodes into non-overlapping modules, such that each brain region gets assigned to one community and one community only [23, 24]. Applying the same algorithm to eFC results in a non-overlapping partition of edges into communities. However, when edges are mapped back to their respective nodes, non-overlapping edge partitions yield overlapping nodal partitions, such that a single node can be associated with multiple communities [25, 26].

In a previous paper, we characterized the basic properties of eFC, including its modular structure [21]. However, the relationship between modules derived from eFC and brain systems derived from nFC remains unclear. Are the edges that link brain systems to one another homogeneous in terms of their edge community assignments? Or are brain systems linked to one another *via* diverse assemblies of edges that comprise several distinct edge communities [27, 28]? Addressing these questions would add clarity to our understanding of how the brain’s modular structure helps support cognition.

Here we investigate this relationship in greater detail with eFC estimated using Midnight Scan Club data [29, 30]. First, we derive edge communities and show that individual brain regions participate in many different communities. Next, we investigate how these communities are distributed within and between traditionally-defined brain systems. We demonstrate that all systems are linked to one another *via* multiple distinct edge communities. Focusing on the configuration of edge communities within brain systems, we use a data-driven community detection algorithm to uncover their multi-scale organization [31], demonstrating that higher-order cognitive systems exhibit more complex communities compared to sensorimotor systems. We then apply the same clustering algorithm to data from the entire cerebral cortex, identifying a novel cluster structure that deviates, systematically, from previously described brain systems. Finally, we investigate edge community structure at the level of individual subjects. We show that edge community structure exhibits remarkable idiosyncrasies, which are driven by the personalization of edge communities outside of sensorimotor cortices. The results presented here offer pathways for future studies aimed at relating features of edge-centric networks to individual differences in behavior and cognition.

## RESULTS

In this section, we present analyses of eFC estimated using resting-state data from the Midnight Scan Club (MSC). Complete details of MRI acquisition, preprocessing pipelines, and network construction can be found in **Materials and Methods**.

### Edge communities reveal overlapping network structure

Many studies have shown that the brain exhibits modular structure, meaning that its elements can be partitioned into cohesive clusters called *communities* or *modules* [10, 11, 32, 33]. Modules are usually defined to be internally dense and non-overlapping (with some notable exceptions [34–37]), such that nodes are assigned to one module only and that nodes tend to be strongly connected to other nodes in their own module and weakly connected to nodes in other modules. Recently, we developed a novel edge-centric representation of brain networks, which we used to cluster network edges, resulting in overlapping nodal communities. Here, we replicate those findings using data from the Midnight Scan Club. We show that community overlap varies across cerebral cortex and canonical brain systems [38]. These observations motivate a further exploration of the relationship of brain systems and edge communities.

We first derived group-representative edge communities. To do so, we estimated edge time series for all 100 resting-state scans in the dataset (10 subjects; 10 scans each), concatenated these data, and used a two-stage clustering algorithm to generate 250 estimates of communities before synthesizing these results into consensus edge communities. These communities can be visualized in several different ways. First, because the clustering algorithm operates at the level of edges, we can visualize edge communities in matrix form, by labeling the edge between nodes *i* and *j* according to its edge community assignment (Fig. 1*b*). Here, each color corresponds to a different edge community (as in our previous paper, we show results with the number of communities fixed at *k* = 10; see Fig. S1 for edge communities detected at other *k*). A second strategy for visualizing edge communities is to calculate for each node the fraction of its edges that belong to a given community. This procedure is especially useful, as it allows us to describe edge communities more intuitively in terms of brain regions and systems (Fig. 1*c*). This also allows us to visualize the topography of edge communities in anatomical spaces, by projecting regional participation in edge communities onto brain surfaces (Fig. 1*d*).

**FIG. 1.**
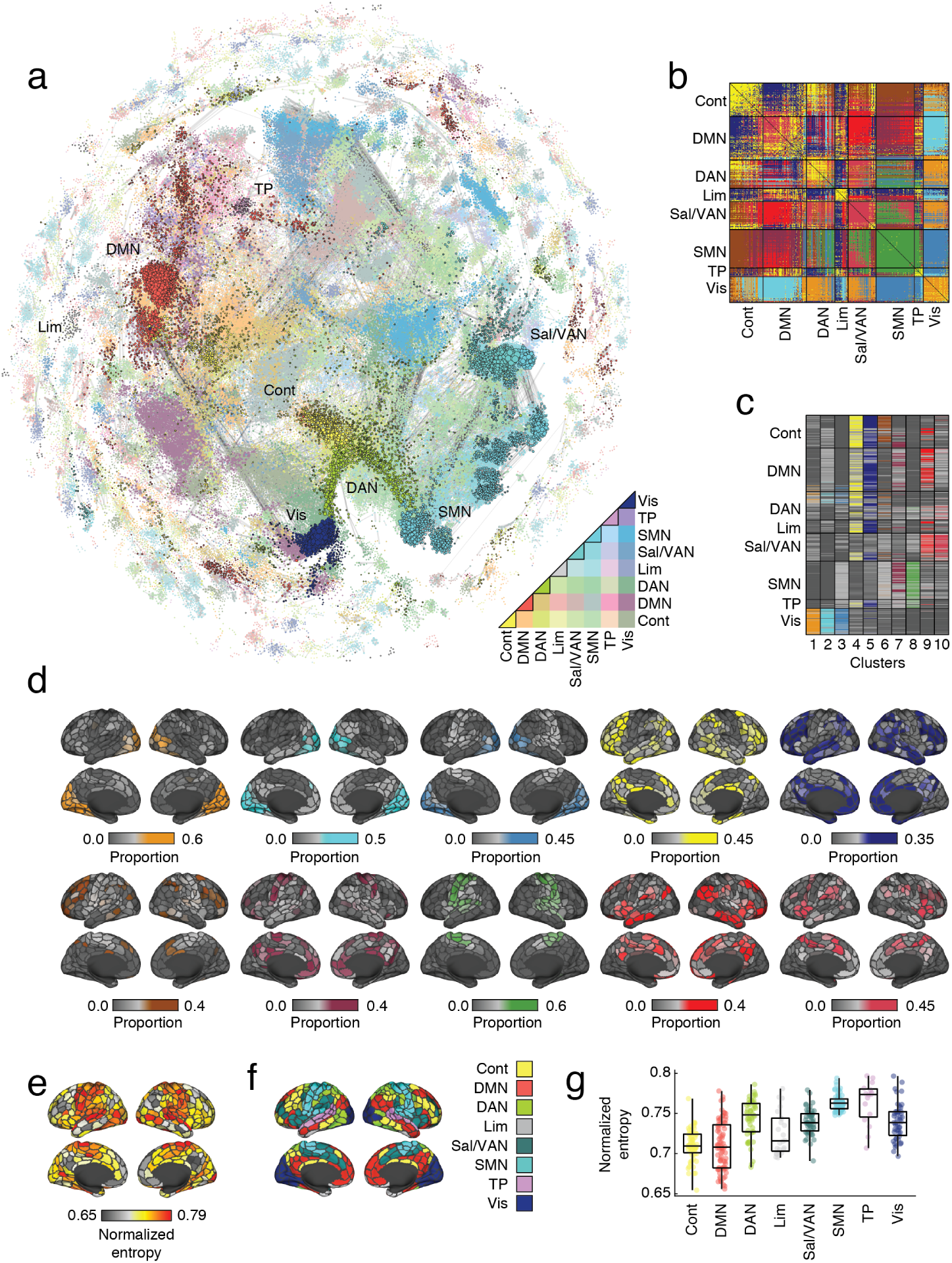
Edge functional connectivity. (*a*) Force-directed layout of edge functional connectivity (eFC). Each point represents an individual edge, colored according to the brain systems to which the edge’s stub nodes belong to. (*b*) Edge communities mapped into a node × node matrix. Each color reflects a distinct edge community. (*c*) Edge communities mapped back to individual nodes. In this plot, rows and columns represent nodes and communities, respectively. Within each column, colors indicate the fraction of a node’s edges that are associated with the corresponding edge community. We can project the columns of this matrix onto the cortical surface. In panel *d* we show projections for each of the *k* = 10 edge communities. From edge communities, we can also calculate the normalized entropy for each node – a measure of community overlap, In panel *e* we show projections of those overlap scores onto the cortical surface. (*f* and *g*) We can then aggregate, entropy (overlap) scores according to brain systems. As in our previous paper, we find that overlap is greatest in primary sensory and attentional systems and lowest in association cortices.

Following our previous paper, we then calculated the level of community overlap for a given brain region as normalized entropy, where values close to 0 indicate that a brain regions’ edges are concentrated among a small number of communities, while values close to 1 indicate that edges are uniformly distributed over communities (Fig. 1*e*). Notably, we found that there were no regions with entropies near zero, in agreement with the observation from our previous paper that brains exhibit “pervasive overlap.” Nonetheless, the community overlap measure exhibited cortical specificity. Again, in agreement with our previous paper, we found that the greatest levels of overlap were concentrated in primary sensory and attentional networks (Fig. 1*f,g*). This observation indicates that the connections associated with brain regions in those systems are involved in many different edge communities. In contrast, heteromodal association cortices, which include control, default mode, and limbic networks, exhibited the lowest levels of overlap.

Collectively, these results recapitulate the main findings from our previous paper [21], and extend them to an increasingly fine-grained parcellation [38]. More practically, the fact that we could obtain qualitatively similar community structure and overlap by clustering edge time series, which are more computationally tractable than the edge connectivity matrix, makes it possible to perform additional complex analyses in the future. In summary, these findings are in line with our earlier report [21] and provide a baseline for the following extension of the edge connectivity framework.

### System-level complexity of edge community structure

An edge community is a collection of edges – pairs of nodes – whose co-fluctuations follows a similar time course. How are these communities distributed within and between canonical brain systems [10, 11, 38]? Are some brain systems linked to one another *via* many communities? Are others linked by few? Here, we address these questions by considering edge community templates – binarized maps of edge communities – which we aggregate into descriptions of system-level interactions. In general, we find additional evidence of “pervasive overlap” [25], such that virtually all pairs of systems are linked to one another by at least two edge communities. We also find that, internally, sensorimotor systems are spanned by relatively few edge communities compared to higher-order heteromodal systems.

We first mapped edge community labels into a node-by-node matrix (Fig. 2*a*) and, for each edge community, extracted its template pattern (Fig. 2*b*), in which edges belonging to that community were assigned a value of 1 while all other edges were set equal to 0. We aggregated the nonzero elements in each template by cognitive systems, counting the fraction of the edges within or between those systems that belonged to a given edge community (Fig. 2*c*). These system-by-system maps quantified the extent to which systems were linked by a given edge community (Fig. 2*d*).

**FIG. 2.**
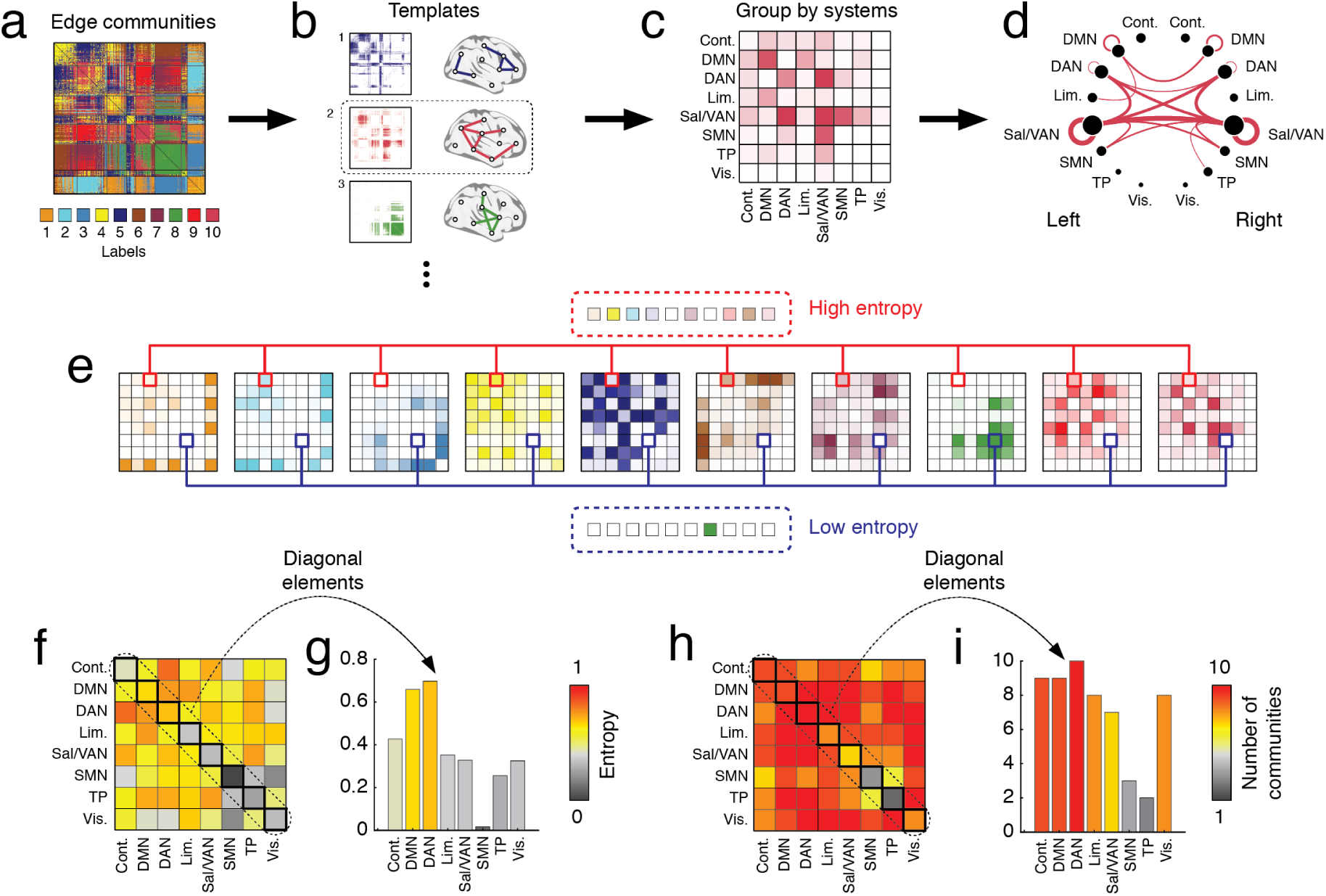
Edge community templates reveal system-dependent organization. (*a*) Edge communities mapped into a node × node matrix. (*b*) We generated community templates, where each community is represented as a binary matrix with edges assigned a value of 1 or 0 depending upon whether they were included in that community. (*c*) We aggregated template edges by brain systems, counting the number of edges that fell within or between eight canonical brain networks. (*d*) Each template describes the fraction of inter- and intra-system interactions mediated by a given edge community (here, we split systems into their left- and right-hemisphere components for visualization only). (*e*) We can use these templates to identify brain systems linked to one another by edges assigned to many or few edge communities (high or low entropy). (*f*) We calculated the entropies for all pairs of brain systems. (*g*) If we consider only within-system edges, we find that heteromodal association cortex tends to have greater entropy and participate in a greater number of discrete edge communities than primary sensory systems (panels *h* and *i*).

Using the system-by-system maps, we estimated the entropy associated with all pairs of systems (Fig. 2*e*). Intuitively, if the edges between those systems belonged to a diverse set of edge communities, then the entropy score was high. On the other hand, if the edges belonged to relatively few communities, then the entropy was low. Interestingly, we found that the highest levels of entropy were associated with connections between the dorsal attention and cognitive control networks, while the lowest were associated with the within-system connections of the somatomotor network (Fig. 2*f*). When considering just the internal edges of brain systems, we found that default mode and dorsal attention had the highest levels of entropy while, somatomotor, temporparietal, and the visual network were among the lowest. Similar patterns were when considering the number of distinct edge communities observed in within- and between-system blocks (Fig. 2*h,i*).

Finally, we investigated the structure of each edge community in greater detail, focusing on the specific brain systems that it linked. Broadly, edge communities could be sub-divided into two groups: “cohesive” communities that included disproportionately many within system edges and “bridge” communities, comprised mostly of edges that fell between brain systems (Fig. 3*a*). We further sub-classified “bridge” communities based on the systems that they linked: “association” bridges linked heteromodal systems (control, default mode, dorsal attention, limbic, salience/ventral attention, and temporoparietal systems) to one another, while “processing” bridges linked heteromodal and unimodal systems (somatomotor and visual) to each other (Fig. 3*b*). As expected, we found that cohesive communities contained a greater proportion of within-system edges than bridge communities (Fig. 3*b*; *p* < 0.05; t-test). We also found that association communities contained a greater proportion of edges linking heteromodal systems to one another compared to processing communities (Fig. 3*c*; *p* < 0.05; t-test) while processing communities contained a greater proportion of heteromodal to unimodal edges (Fig. 3*d*; *p* < 0.05; t-test). We show the full ontology of edge communities in Fig. 3*e*. We find similar results with different numbers of edge communities (Fig. S2).

**FIG. 3.**
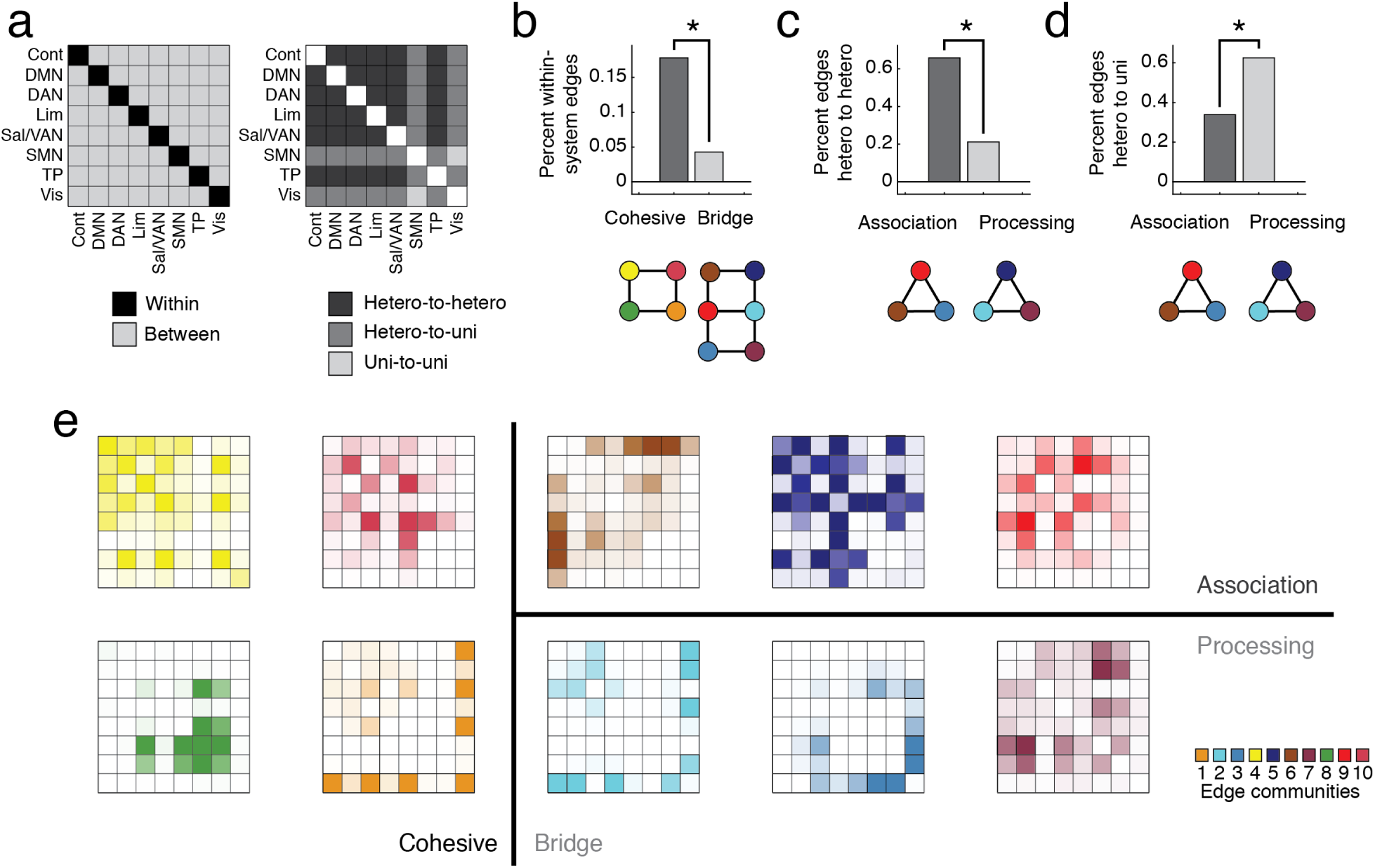
Categorization of edge communities. (*a*) Each edge community was classified as a “cohesive” or “bridge” community according to whether edges belonging to that community fell within or between brain systems, respectively. We further sub-classified bridge communities according to whether the edges linked heteromodal systems (control, default mode, dorsal attention, salience/ventral attention, limbic, temporoparietal) to other heteromodal systems or to sensorimotor systems (somatomotor, visual). We referred to these two sub-categories as “association” and “processing” communities, respectively. (*b*) As expected, we found that cohesive communities included a greater proportion within-system edges compared to bridge communities. Similarly, association communities had a greater proportion of edges linking heteromodal systems to other heteromodal systems (panel *c*) while processing communities exhibited a greater proportion of heteromodal to unimodal edges (panel *d*). In panel *e*, we show the ten edge communities divided into their respective classes. The vertical line divides “cohesive” from “bridge” communities while the horizontal line divides “association” from “processing”. The outlines (black, green, red) are used to help identify system pairs responsible for that edge community’s classification.

Collectively, these findings suggest that the brain’s edge community structure is pervasively overlapping, such that all pairs of brain systems are linked to one another *via* multiple edge communities that, in turn, reflect distinct patterns of edge co-fluctuations. Second, these findings further suggest that although all systems interact *via* distinct modes, the number and diversity of modes is system-dependent and that heteromodal systems exhibit a more complex internal structure than sensorimotor systems. Further, the particular configuration of edge communities among brain systems suggests distinct functional classes, with some edge communities positioned to maintain the cohesiveness of systems and others to form links across system boundaries.

### Multi-scale and system-dependent organization of edge community structure

In the previous section, we showed that brain systems can be linked to one another *via* different modes of coupling (edge communities). Notably, we found that the diversity of edge communities within brain systems was highly variable. Here, we investigate the internal structure of brain systems in greater detail. To do so, we introduce the concept of an *edge community profile* and define a measure of similarity for comparing profiles between pairs of regions. Separately for each cognitive system, we generate the interregional similarity matrix among all regions assigned to that system, which we partition using multi-scale modularity maximization [23, 39–41]. We find that the number of distinct sub-communities within each brain system was greatest for higher-order cognitive systems, whereas sensorimotor networks exhibited many fewer sub-communities.

To estimate multi-scale community structure, we leveraged the node-by-node matrix representation of edge communities (Fig. 4*a*) and extracted each region’s edge community profile as the corresponding row (Fig. 4*b*). To measure the similarity between two region’s profiles, we simply measured the fraction of their elements assigned to the same edge community. Repeating this process for all pairs of brain regions generated a node-by-node similarity matrix (Fig. 4*c*). Considering separately the within-system elements for each system, we found that visual and sensorimotor systems exhibited significantly greater levels of similarity compared to the other brain systems (permutation test, *p* < 10^−3^; Fig. 4*d,e*). We found similar results with different numbers of edge communities (Fig. S3).

**FIG. 4.**
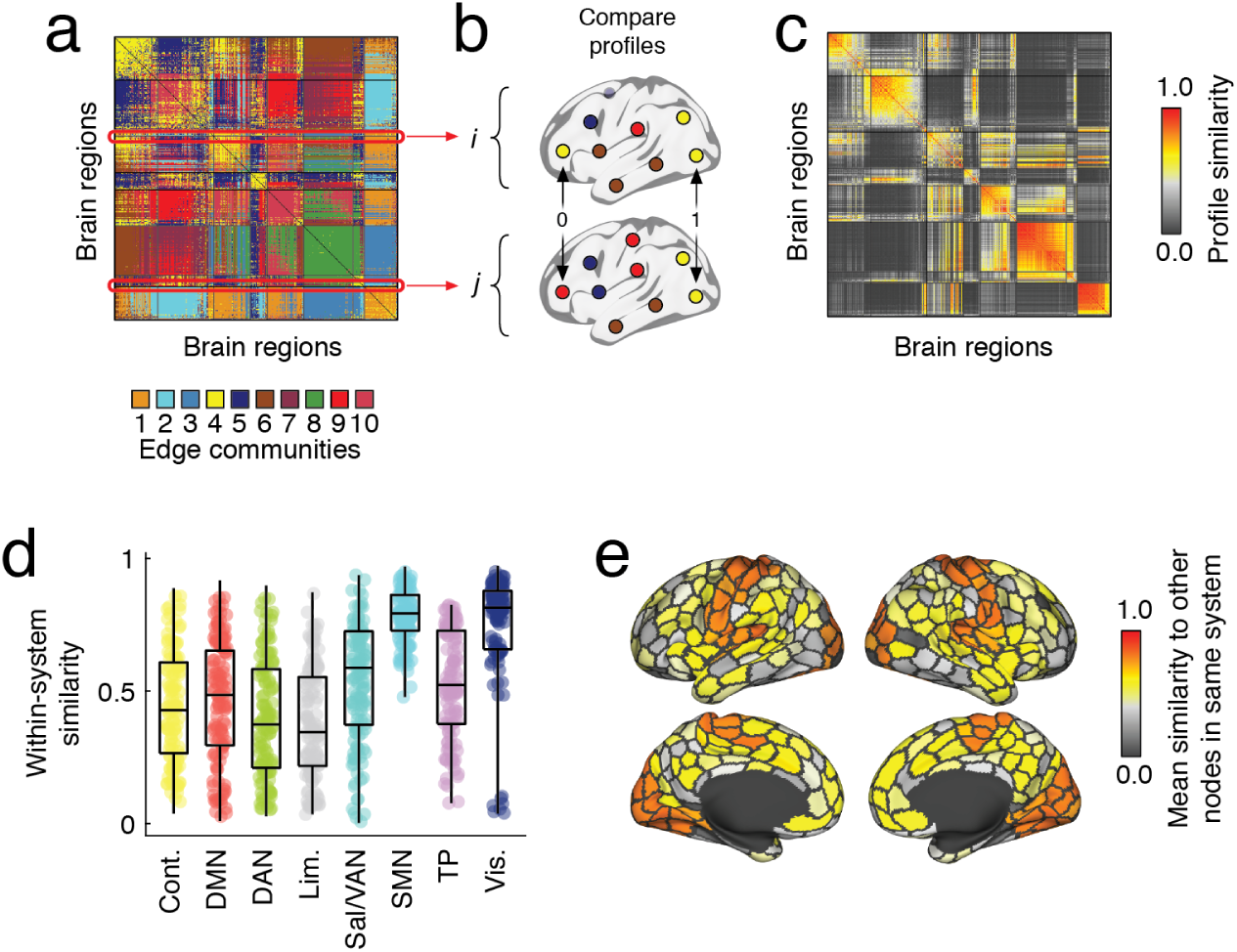
Edge community similarity. (*a*) Edge communities reshaped into a node × node matrix. (*b*) We can treat the columns and rows of this matrix as “profiles” for different regions, and compare nodes’ community labels to measure the similarity of two profiles with respect to one another. Repeating this process for all pairs of nodes results in a similarity matrix (*c*). The average similarity between nodes within each brain system is highly variable. We find that control networks exhibit low levels of overlap and are composed of nodes with heterogeneous edge community profiles (see panel *d*; individual points in this panel correspond to the edge community similarity for pairs of brain regions within a given system). In contrast, we find that sensorimotor networks (visual + somatomotor) exhibit high levels of overlap, but are composed of nodes with homogeneous edge community profiles. We can also visualize the heterogeneity of each system by projecting their mean internal similarity onto the cortex (*e*). Note that similarity is greatest for visual and somatomotor systems.

Next, we clustered the within-system similarity matrix for each system (Fig. 5*a*). This procedure entailed extracting the set of within-system similarity values and, using a variant of modularity maximization [23], estimating clusters across a range of topological scales (by varying the value of a structural resolution parameter, *γ*, over the interval [0, 1] in increments of 0.002 [39]). We then grouped together clusters estimated using similar parameter values and, from these estimates, extracted consensus clusters [42]. We repeated this procedure for multiple topological scales (resolution parameters); here we focus on the range 0.4 < *γ* < 0.5.

**FIG. 5.**
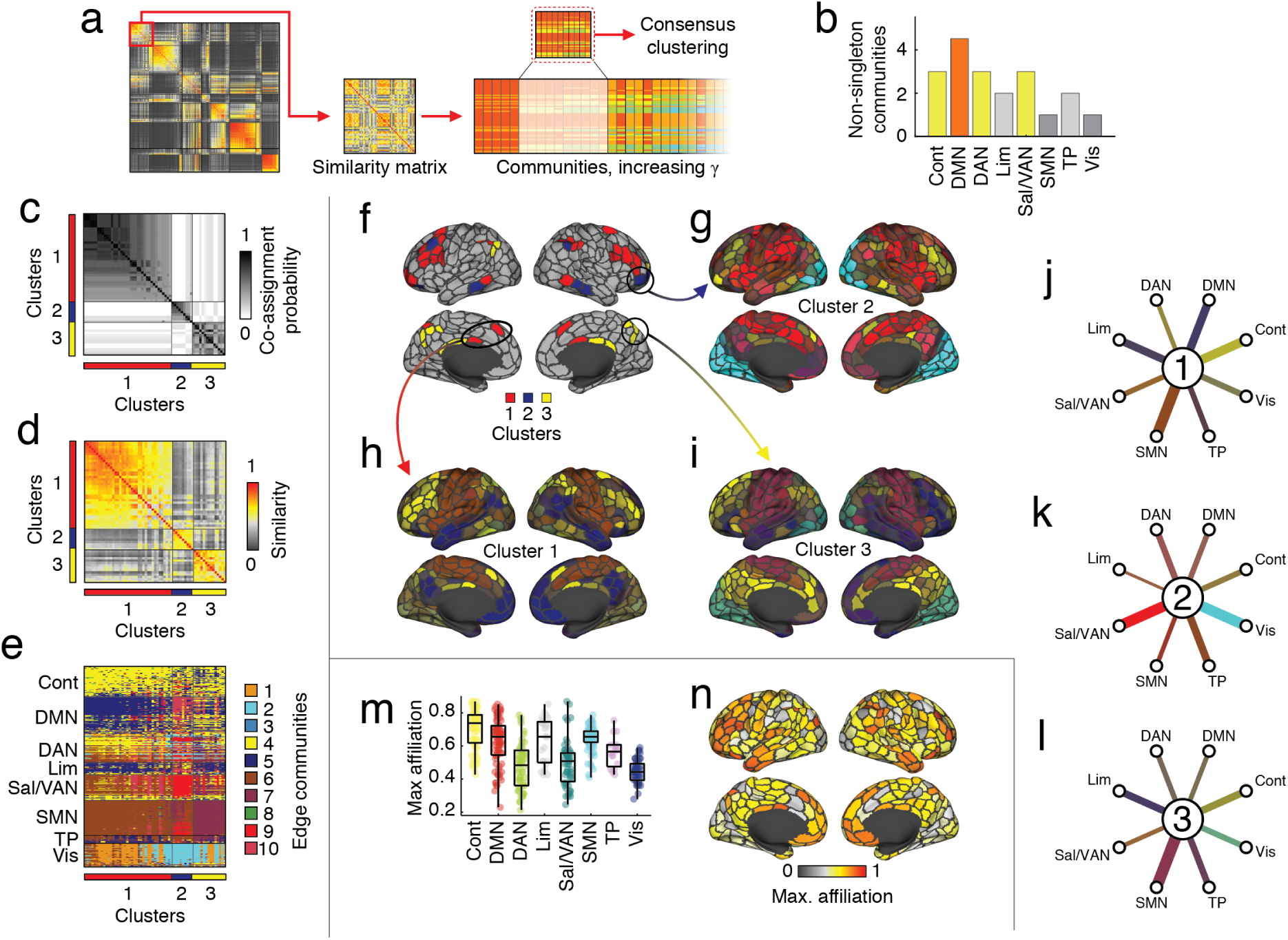
Cluster structure of edge communities. (*a*) Pipeline for estimating system-specific multi-resolution clusters. (*b*) We found that the systems with the greatest number of communities included control, default mode, and both dorsal and ventral attention networks. In panels *c* - *n*, we focus on the control network specifically. (*c*) Co-assignment matrix ordered by consensus communities. (*d*) The within-system similarity matrix ordered according to the three-cluster solution. (*e*) Edge community profiles ordered according to clusters. Topographic representation of consensus clusters (*f*) and cluster centroids (*g* -*i*). In each centroid plot, nodes are colored according to the mode of their edge community assignments emanating from the control network. The brightness of nodes indicates “cluster homogeneity.” (*j* - *l*) Hub and spoke plots for each centroid revealing the dominant edge community linking centroids to brain systems. Maximum affiliation of control nodes to any of the 10 edge communities aggregated by brain system (*m*) and displayed topographically (*n*).

In general, we found that the number of detected clusters was greatest in higher-order systems compared to (Fig. 5*b*). We find similar results at other ranges of *γ* (see Fig. S4). Here, we focus on the control network (we show results for other brain systems in Fig. S5 and comparisons with other reported sub-divisions in Fig. S6), which the clustering algorithm partitioned into three clusters (Fig. 5*c*). Each cluster was, internally, homogeneous (Fig. 5*d*) and was comprised of regions with distinct edge community profiles (Fig. 5*e*). We show these profiles in greater detail in Fig. 5*f* -*i*.

Intuitively, we can think of these profiles as delineating different patterns by which the activity of regions in the control network and the rest of the brain co-fluctuates. To map these patterns back to brain systems, we calculated the dominant edge community linking each of the three control clusters to the eight canonical systems. We depict these cluster-to-system links as hub and spoke diagrams in Fig. 5*j* -*l*. At the center of each diagram is a hub that represents the set of control regions assigned to that cluster. Those regions are connected to each system by spokes colored according to dominant edge community. For instance, edges from control regions in cluster 1 to regions belonging to the salience/ventral attention system tend to belong to the red edge community, while edges linking that cluster to the visual network tend to be cyan edge community.

Importantly, while this analysis suggests that there exists distinct modes of coordination between control regions and the rest of the brain, there are also some patterns of edge communities shared across the multiple clusters. Specifically, we found that nodes in the control network tend to be linked to one another *via* the same edge community (Figure. 5*m*). On the other hand, the edge community assignments of nodes in the control network to dorsal attention, salience/ventral attention, and visual networks are all highly variable.

Taken together, these findings indicate that the internal structure and complexity of edge communities varies across systems. Building on observations from the previous section and in agreement with the extant literature, we find that the the greatest level of complexity are located in the higher-order, heteromodal brain systems, which are associated with a range of cognitive domains. Our findings suggest that their polyfunctionality may be engendered by the diversity of edge communities profiles.

### Uncovering whole-brain communities from edge community profiles

In the previous section, we used a multi-resolution community detection algorithm to uncover the cluster structure of edge communities within specific brain systems. Although these analyses revealed differences from one system to another, they prevented us from discovering patterns in edge communities at the whole-brain level. For instance, if two nodes had identical edge community profiles but were assigned to different systems, the previous analyses would be incapable of grouping them together into the same cluster. To address these limitations, we used the same algorithm as in the previous section to uncover multi-scale community structure using whole-brain data. We found that clusters derived from edge communities largely approximated known cognitive systems. However, we also uncovered subtle yet systematic differences between nodes’ assigned clusters and their canonical system labels.

We applied a multi-resolution consensus clustering algorithm to partition the cerebral cortex into non-overlapping communities of different sizes (Fig. 6*a*). Here, we focus on an intermediate scale that resulted in seven large clusters and multiple small clusters (which we group into a separate cluster for convenience) (Fig. 6*b*). We note that the larger clusters tended to be stable across the full range of *γ* values. At an intermediate level (0.4 < *γ* < 0.5), nodes assigned to the detected clusters were similar to one another (Fig. 6*c*), resulting in homogeneous edge community profiles (Fig. 6*d*; we show partitions derived at other resolutions in Fig. S7).

**FIG. 6.**
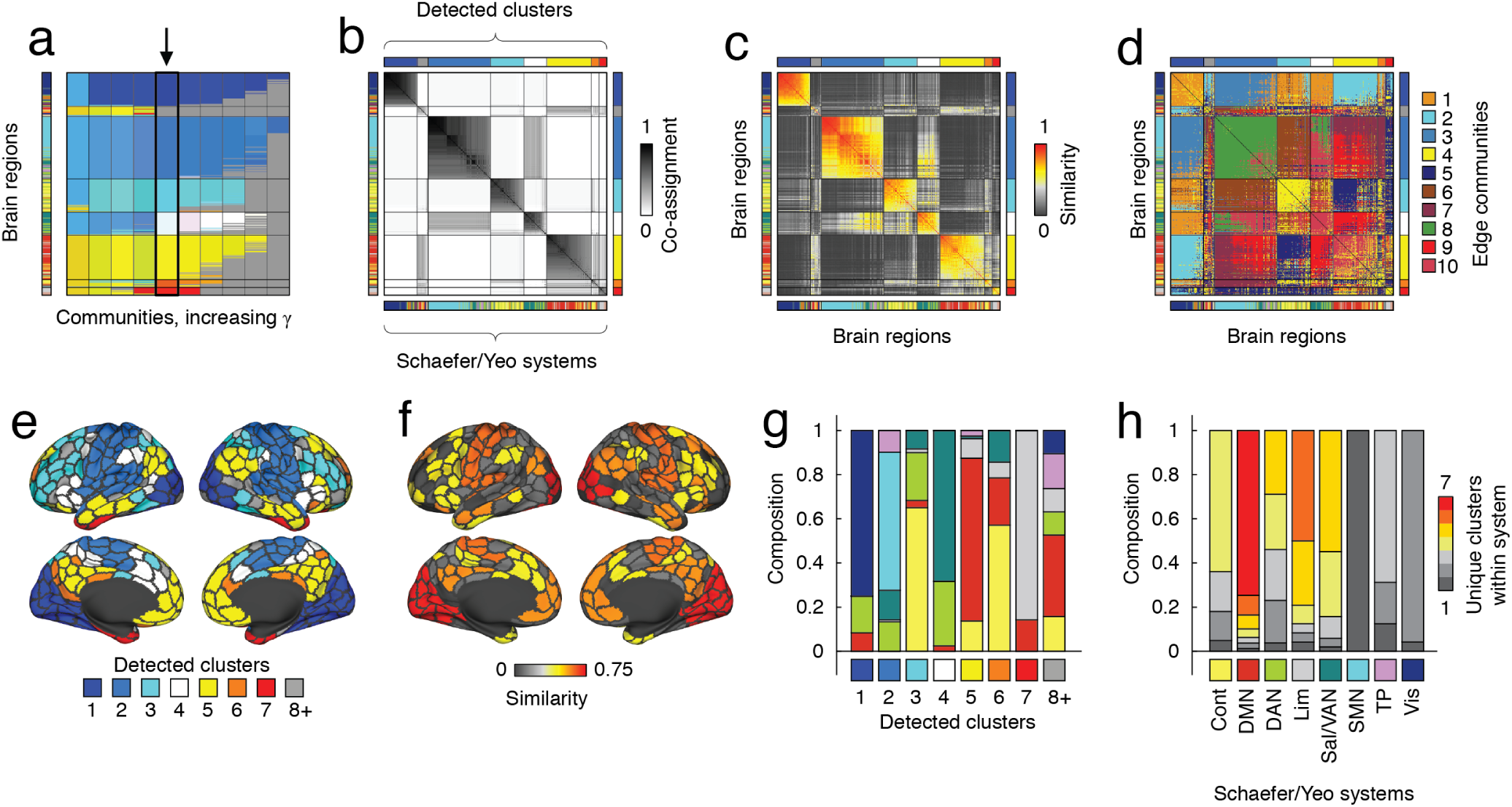
Multi-resolution cluster structure of cortical edge communities. (*a*) Nodes’ cluster assignments arranged in order from coarsest to finest levels. In our analysis we focus on a level where there exists seven large clusters comprised of > 20 nodes each (black arrow). Note that the nodes are ordered by detected clusters and not brain systems. The bar on the *x* -axis depicts brain system labels in the same ordering. (*b*) Interregional cluster co-assignment probabilities, (*c*) whole-brain edge community similarities, and (*d*) edge community labels in matrix form, ordered by detected clusters. (*e*) Clusters mapped onto the cortical surface. Small clusters are collapsed into a single label (gray). (*f*) Regional similarity of detected clusters and canonical brain systems. (*g*) System composition of detected clusters. (*h*) Cluster composition of brain systems. Color indicates the number of unique clusters within a system.

Broadly, the detected clusters were similar to known brain systems (Fig. 6*e*). To assess this correspondence more directly, we computed the similarity (Jaccard index) of each region’s assigned cluster and system. Overall, the visual and somatomotor networks exhibited greater than expected similarity while control, dorsal attention, limbic, and temporparietal networks were more dissimilar than expected (*p*_*adjusted*_ = 0.0016; false discovery rate fixed at 5%; Fig. 6*f*). To better visualize the overlap, we calculated the composition of each cluster in terms of their assigned nodes’ system labels (Fig. 6*g*). We found that clusters 1, and 2 were almost uniformly composed of regions from the visual and somatomotor systems. The other clusters were less homogeneous, and received substantial contributions from multiple brain systems. Interestingly, cluster 8 (which was an aggregate of all the small communities) included relatively few sensorimotor nodes and was composed of regions from control, default mode, attention, limbic, and tempoparietal systems. We performed a similar analysis, grouping the detected clusters by brain systems (Fig. 6*h*). We found that visual and somatomotor systems were composed of relatively few distinct clusters, whereas the other brain systems were composed of nodes from multiple different clusters.

Collectively, these results suggest that the similarity of regions’ edge community profiles is largely aligned with the brain’s known system-level organization. However, we also find that differences between the two sets of labels follow a distinct pattern. Misalignment tends to involve regions typically assigned to heteromodal systems.

### Edge community structure is subject-specific

To this point, all analyses have focused on relating brain systems to edge communities using pooled, group-representative data. These analyses uncovered shared relationships, common across a small cohort of individuals. However, there remain several important unresolved questions. For instance, to what extent are edge communities variable across individuals? Are the edge community profiles of some regions and systems differentially variable across individuals? Does variability of those features reflect meaningful, subject-specific traits? Here, we address these questions by detecting and comparing edge communities within and between subjects and scans.

To address these questions, we performed three separate analyses. First, for each subject we concatenated their scans and estimated their subject-specific consensus edge communities (Fig. 7*a*). Subjects’ edge communities were more similar to group representative partition than expected by chance (permutation test; *p* < 10^−3^).

**FIG. 7.**
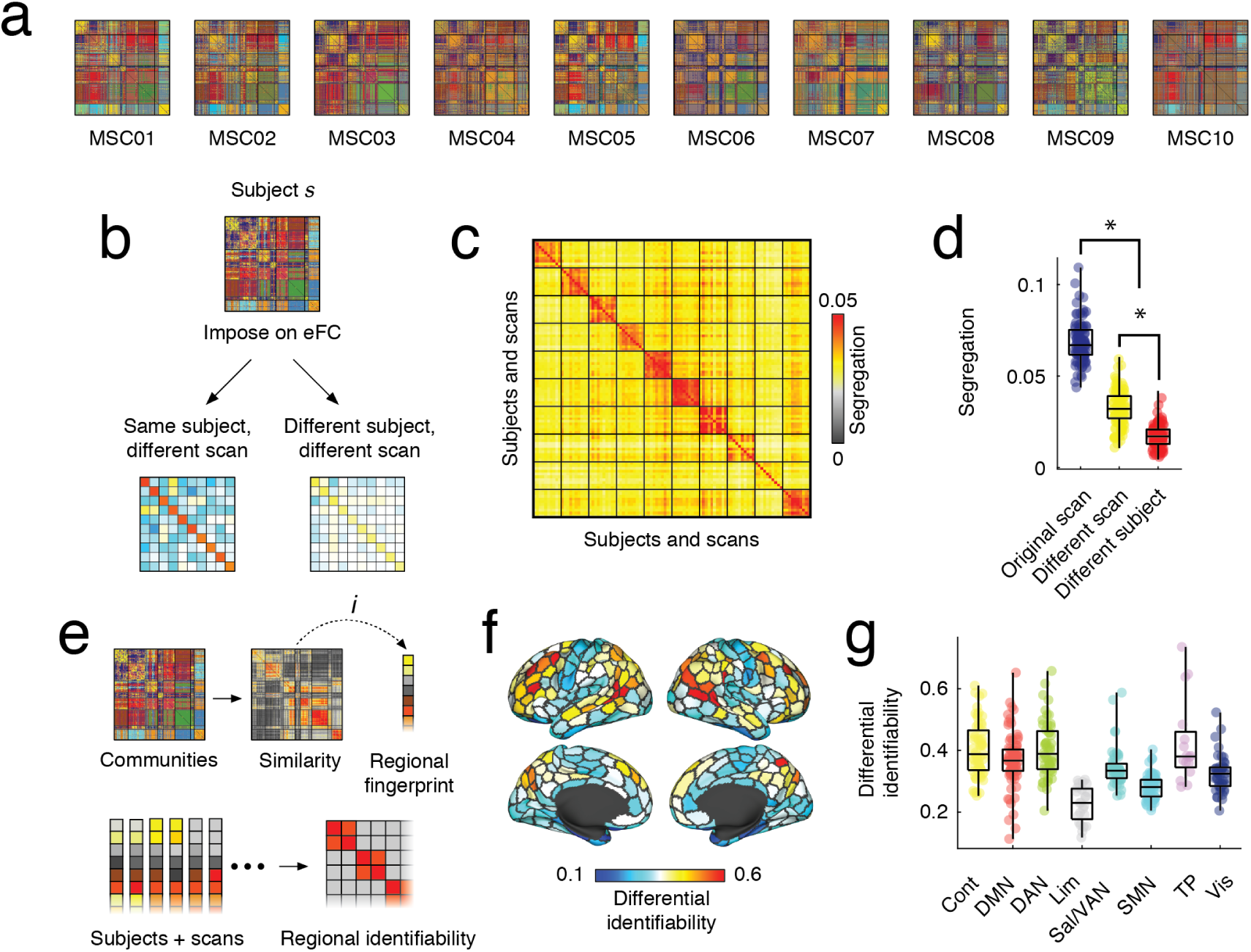
Personalization of edge community structure. (*a*) Subject-representative edge communities (estimated using ten scans). (*b*) Pipeline for segregation estimation. We derive edge communities for each subject and each scan and impose those communities onto eFC estimated from all other scans and subjects. Segregation is measured as the mean within-community eFC minus the mean between-community eFC. (*c*) Scan-by-scan matrix of segregation scores; rows represent the subject and scan from which edge communities were estimated and columns represent the subject and scan onto which those communities were imposed. The brightness of cells represents the level of segregation. (*d*) Comparing segregation scores within and between scans/subjects. (*e*) Pipeline for calculating regional differential identifiability. (*f*) Topographic representation of regional differential identifiability scores. (*g*) Regional differential identifiability scores aggregated by brain systems.

Visual inspection revealed that edge communities were heterogeneous across subjects, suggesting that edge communities might capture idiosyncratic and subject-specific variation. To test this hypothesis, we estimated edge communities for each subject and each scan (Fig. 7*b*). If edge communities were unique to individual subjects, then we would expect that imposing them on another scan from the same subject would result in segregated edge communities (strong internal eFC; weak external eFC). On the other hand, imposing those edge partitions onto eFC from a different individual would result in reduced segregation. We tested this hypothesis by systematically imposing each of the 100 edge partitions onto eFC estimated from 10 subjects and their 10 scans (100 scans in total) and calculated the “segregation” score as a the mean within-community eFC minus the mean between-community eFC (Fig. 7*c*). As expected, we found that segregation was greatest when we imposed a partition back on the eFC used to estimate those edge communities in the first place (Fig. 7*d*). Interestingly, we also found the segregation was greater when we imposed edge communities on eFC estimated from the same individual than on eFC estimated from other individuals (Fig. 7*d*). These observations suggest that edge communities capture meaningful, subject-specific patterns of edge-edge interactions.

These analyses, however, did not reveal what parts of the brain make subjects identifiable. Here, we address this question by estimating the differential identifiability associated with the edge community structure of every brain region. Specifically, for a given scan and subject, we can generate a vector region *i*’s similarity with respect to all *j* ≠ *i* (Fig. 7*e*). We can then extract analogous vectors from that subject’s other scans and from all subjects’ and their respective scans. Calculating the matrix of pairwise correlations, we compute the differential identifiability as the mean within-subject similarity minus the mean between-subject similarity. We then repeat this procedure for all regions.

This procedure generates a score for every brain region that describes, on average, how personalized and idiosyncratic its edge communities are. In Fig. 7*f*, we show those scores projected onto the cortical surface. Interestingly, we find considerable variability across the cortex in terms of identifiability, with regions in the control network, along with temporparietal and dorsal attention networks performing particularly well (Fig. 7*g*). We find similar results using different numbers of communities (see Fig. S8).

In summary, these results further implicate the control network, along with other areas in attentional and temporparietal networks, as key drivers of individuality in edge communities. Our work builds on a previously established quantitative framework for tracking identifiable features of brain imaging and network data [43], and extends this framework using edge connectivity data. In doing so, we rely on a mapping of edge communities back into a node-centric framework, thereby improving their interpretability.

Overall, these findings suggest that edge communities are highly personalized, and that this personalization can be linked to the variability of edge communities associated with many different systems in general, but in particular the cognitive control network. These observations agree with other recent studies reporting that control networks carry personalized information about subjects [44]. In summary, our findings underscore the inter-subject variability of the brain’s community and system-level architecture, complementing companion analyses of MSC data using node-centric models of connectivity [29, 30].

## DISCUSSION

In this paper, we investigated the configuration of edge communities across canonical brain systems. We found that all pairs of systems were linked to one another by at least two edge communities and that the exact number and diversity of such links varied by system. Focusing only on within-system edges, we found that the variability and diversity of edge communities comprising higher order cognitive systems was greater than that of sensorimotor systems. We then used a data-driven clustering algorithm to partition brain regions in each brain system into multi-scale communities according to the similarity of their edge community profiles. We found that the number of detected communities is greatest in heteromodal systems and lowest in sensorimotor systems. Repeating this analysis using data from the complete cerebral cortex, we discovered that, overall, the detected clusters resembled known brain systems. However, there were also systematic discrepancies between system labels and the detected clusters, revealing incongruity between clusters derived from traditional nFC and those derived from eFC. Finally, we show that edge community structure is subject-specific and reproducible across multiple scans of the same individual. This personalization is driven by the edge community assignments of nodes located in control, default mode, dorsal attention, and temporparietal networks.

### Pervasive overlap and multiplexity

Many studies have partitioned brain regions based on their functional connections, revealing a surprisingly consistent set of communities that align well with activation patterns and well-known brain systems [10, 11, 13, 45]. These observations suggest that assortative and segregated communities may play an important role in the emergence of functional specialization. Here, rather than focus on partitions of brain regions into communities, we leveraged a recently-proposed edge-centric network model to partition connections into communities [21, 22]. The resulting edge communities delineate groups of functional connections whose valence and amplitude co-fluctuate with one another over time. We speculate that these co-fluctuation patterns may correspond to distinct modes of interregional communication.

A key question, then, was whether edge communities were aligned with the boundaries of traditionally-defined brain systems. That is, if we were to examine the complete set of connections between regions in systems *A* and *B*, would those connections co-fluctuate uniformly and be assigned to a single edge community, or would they be composed of several distinct patterns of co-fluctuation? Phrased alternatively and in line with the hypothesis that co-fluctuating edges reflect distinct modes of inter-regional communication – do systems communicate with one another through a single homogeneous mode or do they communicate in parallel *via* a series of multiplexed channels? Here, we addressed this question by counting the number and distribution of edge communities linking pairs of systems. In all cases, systems were linked by multiple edge communities, although the number and diversity varied considerably across system pairs. These observations suggest that the brain exists in a state of “pervasive overlap” [21, 25], where regions and systems throughout the brain are linked to one another through multiple edge communities.

Our findings have important implications for understanding brain function. In most studies, brain regions are assigned to non-overlapping communities with distinct functional profiles [10, 11, 46]. Polyfunctionality emerges from this caricature in the form of a small subset of brain regions whose connectivity patterns span system boundaries [19, 47]. On the other hand, we find that all brain regions participate in many communities and the functional connections bridging brain systems are associated with a plurality set of community labels. These observations suggest that overlapping function may be a key organizing principle of brain networks, and a rule rather than an exception.

Why, then, do we observe multiplexed, overlapping community structure in the brain? Why are the same brain systems linked by dissimilar patterns of co-fluctuation? One obvious possibility is that the current system ontology does not fully capture the sub-divisions and fine-scale structure of cortical architecture [48]. That is, edge communities may reveal organization that is obscured by or inaccessible using node-centric network models. Another possibility is that edge communities reflect a form of functional robustness and redundancy [49]. That is, by communicating across multiple “channels”, brain systems reduce the likelihood that damage to any one channel would result in a complete disruption of communication and brain function [50–53]. Future work is necessary to clarify the precise functional roles of multiplexed and overlapping communities.

### Heterogeneity and system-specificity of edge community profiles

Here, we examined edge communities from the perspective of brain regions by defining edge community “profiles”. Focusing on profiles, we were able to map edge communities from an unfamiliar and large *m*-dimensional edge space back into a *n*-dimensional node space. By studying the similarity of regions’ profiles to one another, we were able to characterize the diversity of edge communities among regions that make up traditional brain systems. Using this approach, we generated region-by-region similarity matrices for every system and clustered them using a multi-resolution algorithm.

Interestingly, the internal structure of edge community profiles varied across brain systems, with the regions in sensorimotor systems exhibiting highly similar edge community profiles and regions in higher order, heteromodal systems exhibiting greater variation. These observations agree with current theories of cortical organization and function. In terms of node-centric community structure, sensorimotor systems are among the most functionally segregated [10, 54] and occupy opposite positions along smoothly varying functional gradients [55].

The same analysis pipeline was applied to similarity matrices constructed using edge community profiles from the entire cerebral cortex. Specifically, the detected communities resembled known system-level divisions of cortex [38]. We found that regions associated with higher-order brain systems were more likely to fragment and form small (sometimes singleton) clusters with distinct edge community profiles. Importantly, the detected clusters were inhomogeneous and contained regions associated with multiple brain systems. Collectively, these findings suggest that edge communities give rise to distinct regional profiles that are organized into clusters that span traditional system-level boundaries.

### Personalization of edge community structure

Most of this report focused on edge community structure using composite edge time series assembled from multiple subjects. Although analysis of group-representative data can uncover patterns of eFC shared across many individuals, it is poorly suited for uncovering personalized and idiosyncratic features of eFC, which are key elements necessary for biomarker generation [56, 57]. Addressing this limitation, we derived edge communities for the ten individuals in the Midnight Scan Club dataset. We found that subjects’ edge community structure was idiosyncratic, so communities estimated from subject *s* using data from scan *t* did a good job describing edge communities of the same subject on scan *t*′ but a poor job describing edge communities of any other subject. Importantly, these idiosyncrasies arise from the community assignments of edges associated with control, default mode, dorsal attention, and temporoparietal networks.

These observations agree with other recent analysis of MSC data, reporting high levels of personalization in both cortical and subcortical networks [29, 30, 58, 59]. Like similar findings in larger populations [43, 44] our findings implicate heteromodal association cortex as being both highly repeatable across scans of the same subject but maximally dissimilar across individuals. These observations suggest that edge communities, which we interpret as modes of temporally-resolved accounts of ongoing communication between brain regions, are also subject-specific and personalized. We further link the personalization of edge community structure to the assignments of edges associated with higher-order cognitive systems, including control, default mode, dorsal attention, and temporoparietal networks. The findings reported here align with other recent studies suggesting brain network organization is highly individualized [29, 30, 58, 60–62]. Collectively, these observations open up the tantalizing prospect of more targeted and increasingly personalized interventions in the future.

### Future directions

Our work opens up several opportunities for future studies, both methodological and applied. For instance, are inter-individual differences in the number and diversity of edge communities between brain systems related to behavioral, demographic, and clinical variables of interest like a subject’s performance on a cognitively-demanding task [7], their biological age [63], or their neuropsychiatric state [8]? Similarly, future studies should investigate individual differences in the composition and sub-divisions of brain systems. For example, is the complexity and heterogeneity of edge community profiles within subjects’ control networks related to their performance on tasks that require cognitive control, e.g. Stroop or Navon tasks [32, 64]?

Other potentially fruitful opportunities for future studies include exploring subcortical [58] and cerebellar organization [65] with edge communities. These areas were excluded from the present study, but could be investigated in greater detail, yielding new insight into cortical-subcortical interactions [66]. Relatedly, features derived from edge-centric network models, including overlapping communities, could be incorporated into parcellation generation frameworks to create novel cortical parcellations [67].

### Limitations

One overarching limitation surrounding this study concerns the interpretability of eFC. While traditional nFC is now largely accepted within the human neuroimaging community and is frequently interpreted as a measure of interregional communication (although with many caveats [68]), eFC is novel, high-dimensional, and may be difficult to interpret. While this study attempts to form a conceptual bridge between the system-level organization of nFC and edge communities, future work is necessary to clarify, in more precise terms, the relationship between these two constructs.

A second limitation concerns the procedure for estimating edge communities. Here, we use a k-means algorithm that partitions edges into a fixed number of clusters on the basis of their similarity (eFC) with respect to one another. The motivation to use k-means as opposed to other clustering algorithms stems from its computational efficiency and the fact that eFC can be viewed as a distance metric, and can be used by the k-means algorithm to estimate edge communities from edge time series directly. However, there exists a multitude of alternative algorithms that could, in principle, be applied to edge time series or eFC to estimate communities, including the suite of graph clustering algorithms [20, 69], but also time-series decompositions algorithms like independent components analysis (ICA) [70], which has proven especially useful in the analysis of neuroimaging data [71]. Future studies should investigate the effect of clustering algorithm on the character of detected edge communities.

### Conclusion

In summary, detailed analysis of edge functional connectivity and edge communities revealed marked heterogeneity across brain systems and highly reproducible and idiosyncratic patterns within subjects. These findings help establish edge functional connectivity as a useful representational framework and edge communities as measures of potential interest for revealing novel brain-behavior associations and individual differences in brain organization.

## MATERIALS AND METHODS

### Datasets

The Midnight Scan Club (MSC) dataset [30] included rsfMRI from 10 adults (50% female, mean age = 29.1 ± 3.3, age range = 24-34). The study was approved by the Washington University School of Medicine Human Studies Committee and Institutional Review Board and informed consent was obtained from all subjects. Subjects underwent 12 scanning sessions on separate days, each session beginning at midnight. 10 rsfMRI scans per subject were collected with a gradient-echo EPI sequence (run duration = 30 min, TR = 2200 ms, TE = 27 ms, flip angle = 90°, 4 mm isotropic voxel resolution) with eyes open and with eye tracking recording to monitor for prolonged eye closure (to assess drowsiness). Images were collected on a 3T Siemens Trio.

### Image preprocessing

#### MSC functional preprocessing

Functional images in the MSC dataset were pre-processed using *fMRIPrep* 1.3.2 [72], which is based on Nipype 1.1.9 [73]. The following description of *fMRIPrep*’s preprocessing is based on boilerplate distributed with the software covered by a “no rights reserved” (`CC0`) license. Internal operations of *fMRIPrep* use Nilearn 0.5.0 [74], ANTs 2.2.0, FreeSurfer 6.0.1, FSL 5.0.9, and AFNI v16.2.07. For more details about the pipeline, see the section corresponding to workflows in *fMRIPrep*’s documentation.

The T1-weighted (T1w) image was corrected for intensity non-uniformity with `N4BiasFieldCorrection` [75, 76], distributed with ANTs, and used as T1w-reference throughout the workflow. The T1w-reference was then skull-stripped with a Nipype implementation of the `antsBrainExtraction.sh` workflow. Brain surfaces were reconstructed using `recon-all` [77], and the brain mask estimated previously was refined with a custom variation of the method to reconcile ANTs-derived and FreeSurfer-derived segmentations of the cortical graymatter using Mindboggle [78]. Spatial normalization to the *ICBM 152 Nonlinear Asymmetrical template version 2009c* [79] was performed through nonlinear registration with `antsRegistration`, using brain-extracted versions of both T1w volume and template. Brain tissue segmentation of cerebrospinal fluid (CSF), white-matter (WM) and gray-matter (GM) was performed on the brain-extracted T1w using FSL’s `fast` [80].

Functional data was slice time corrected using AFNI’s `3dTshift` and motion corrected using FSL’s `mcflirt` [81]. *Fieldmap-less* distortion correction was performed by co-registering the functional image to the same-subject T1w image with intensity inverted [82] constrained with an average fieldmap template [83], implemented with `antsRegistration`. This was followed by co-registration to the corresponding T1w using boundary-based registration [84] with 9 degrees of freedom. Motion correcting transformations, field distortion correcting warp, BOLD-to-T1w transformation and T1w-to-template (MNI) warp were concatenated and applied in a single step using `antsApplyTransforms` using Lanczos interpolation. Several confounding time-series were calculated based on this preprocessed BOLD: framewise displacement (FD), DVARS and three regionwise global signals. FD and DVARS are calculated for each functional run, both using their implementations in Nipype [85]. The three global signals are extracted within the CSF, the WM, and the whole-brain masks. The resultant `nifti` file for each MSC subject used in this study followed the file naming pattern *_space-T1w_desc-preproc_bold.nii.gz.

### Image quality control

The quality of functional images in the MSC were assessed using *fMRIPrep*’s visual reports and *MRIQC* 0.15.1 [86]. Data was visually inspected for whole brain field of view coverage, signal artifacts, and proper alignment to the corresponding anatomical image.

### Functional and structural networks preprocessing

#### Parcellation preprocessing

A functional parcellation designed to optimize both local gradient and global similarity measures of the fMRI signal [38] (*Schaefer400*) was used to define 400 areas on the cerebral cortex. These nodes are also mapped to the *Yeo* canonical functional networks [11]. For the MSC dataset, a *Schaefer400* parcellation was obtained for each subject using a Gaussian classifier surface atlas [87] (trained on 100 unrelated Human Connectome Project subjects) and FreeSurfer’s `mris ca label` function. These tools utilize the surface registrations computed in the `recon-all` pipeline to transfer a group average atlas to subject space based on individual surface curvature and sulcal patterns. This method rendered a T1w space volume for each subject. For use with functional data, the parcellation was resampled to 2mm T1w space.

#### Functional network preprocessing

Each preprocessed BOLD image was linearly detrended, band-pass filtered (0.008-0.08 Hz) [88], confound regressed and standardized using Nilearn’s `signal.clean`, which removes confounds orthogonally to the temporal filters [89]. The confound regression employed [90] included 6 motion estimates, time series of the mean CSF, mean WM, and mean global signal, the derivatives of these nine regressors, and the squares these 18 terms. Furthermore, a spike regressor was added for each frame exceeding 0.5mm framewise displacement. Following preprocessing and nuisance regression, residual mean BOLD time series at each node were recovered. eFC matrices for each subject were computed and then averaged across subjects, to obtain a representative eFC matrix for each dataset.

### Edge graph construction

Constructing networks from fMRI data (or any neural time series data) requires estimating the statistical dependency between of time series. The magnitude of that dependency is usually interpreted as a measure of how strongly (or weakly) those voxels are parcels are functionally connected to each other. By far the most common measure of statistic dependence is the Pearson correlation coefficient. Let **x**_*i*_ = [*x*_*i*_(1), …, *x*_*i*_(*T*)] and **x**_*j*_ = [*x*_*j*_(1), …, *x*_*j*_(*T*)] be the time series recorded from voxels or parcels *i* and *j*, respectively. We can calculate the correlation of *i* and *j* by first z-scoring each time series, such that 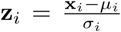, where 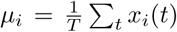 and 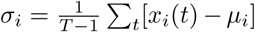 are the time-averaged mean and standard deviation. Then, the correlation of *i* with *j* can be calculated as: 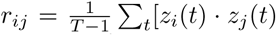. Repeating this procedure for all pairs of parcels results in a node-by-node correlation matrix, i.e. an estimate of FC. If there are *N* nodes, this matrix has dimensions [*N* × *N*].

To estimate *edge*-centric networks, we need to modify the above approach in one small but crucial way. Suppose we have two z-scored parcel time series, **z**_*i*_ and **z**_*j*_. To estimate their correlation we calculate the mean their element-wise product (not exactly the average, because we divide by *T* − 1 rather than *T*). Suppose, instead, that we never calculate the mean and simply stop after calculating the element-wise product. This operation would result in a vector of length *T* whose elements encode the moment-by-moment co-fluctuations magnitude of parcels *i* and *j*. For instance, suppose at time *t*, parcels *i* and *j* simultaneously increased their activity relative to baseline. These increases are encoded in **z**_*i*_ and **z**_*j*_ as positive entries in the *t*th position, so their product is also positive. The same would be true if *i* and *j decreased* their activity simultaneously (because the product of negatives is a positive). On the other hand, if *i* increased while *j* decreased (or *vice versa*), this would manifest as a negative entry. Similarly, if either *i* or *j* increased or decreased while the activity of the other was close to baseline, the corresponding entry would be close to zero.

Accordingly, the vector resulting from the elementwise product of **z**_*i*_ and **z**_*j*_ can be viewed as encoding the magnitude of moment-to-moment co-fluctuations between *i* and *j*. An analogous vector can easily be calculated for every pair of parcels (network nodes), resulting in a set of co-fluctuation (edge) time series. With *N* parcels, this results in 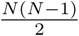 pairs, each of length *T*. From these time series we can estimate the statistical dependency for every pair of edges. We refer to this construct as edge functional connectivity (eFC). Let **c**_*ij*_ = [*z*_*i*_(1) · *z*_*j*_(1), …, *z*_*i*_(*T*) · *z*_*j*_(*T*)] and **c**_*uv*_ = [*z*_*u*_(1)·*z*_*v*_(1), …, *z*_*i*_(*T*)·*z*_*j*_(*T*)] be the time series for edges {*i, j*} and {*u, v*}, respectively. Then we can calculate eFC as:

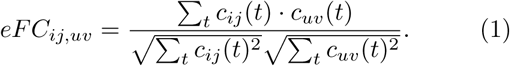

Here, the denominator is necessary to bound eFC to the interval [−1, 1].

### Edge community detection algorithm

In our previous paper we developed a spectral method for clustering eFC matrices [21]. Although this algorithm operated on a reduced rank version of eFC matrices, obtaining these lower rank data required first generating the eFC matrix. In general, eFC matrices are much larger than nFC matrices. This means that they take longer to compute and much more memory. Here, we circumvent this issue by clustering the edge time series directly. A parcellation of the brain into *N* regions results inn *M* = *N* (*N* − 1)*/*2 edges. So rather than generating an *M* × *M* matrix, reducing its dimensionality, and then clustering its low-dimensional representation, we simply cluster the *M* × *T* time series (where *T*) is the number of samples. We use a k-means clustering algorithm where the distance metric is defined as (1 − *eFC*)*/*2. Two perfectly correlated edge time series have a distance of 0 while two orthogonal edge time series would have a distance of 1.

We used this same algorithm to generate estimates of edge communities at the scale of scans, subjects, and cohort. To generate subject-representative communities, we concatenated edge time series from all of a subjects’ scans and clustered the concatenated time series. Similarly, to generate group representative partitions, we concatenated scans from all subjects. At all scales, we repeated the clustering algorithm 250 times.

### Community overlap metrics

The clustering algorithm partitioned edges into non-overlapping clusters. That is, every edge {*i, j*}, where *i, j* ∈ {1, …, *N*}, was assigned to one of *k* clusters. In this list of edges, each node appeared *N* − 1 times (we excluded self-connections). Region *i*’s participation in cluster *c* was calculated as:

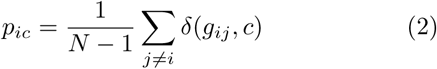

where *g*_*ij*_ ∈ {1, …, *k*} was the cluster assignment of the edge linking nodes *i* and *j* and *δ*(*x, y*) is the Kronecker delta, whose value is 1 if *x* = *y* and zero otherwise.

By definition, ∑_*c*_ *p*_*ic*_ = 1, and we can treat the vector **p**_*i*_ = [*p*_*i*1_, …, *p*_*ik*_] as a probability distribution. The entropy of this distribution measures the extent to which region *i*’s community affiliations are distributed evenly across all communities (high entropy and high overlap) or concentrated within a small number of communities (low entropy and low overlap). We calculate this entropy as:

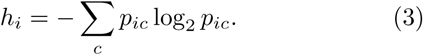

To normalize this measure and bound it to the interval [0, 1], we divide by log2 *k*. We refer to this measure as community entropy and interpret this value as an index of overlap.

### Edge community similarity

When we cluster an eFC matrix, we assign each edge to a single community. These edge communities can be rearranged into the upper triangle of a *N × N* matrix, **X**, whose element *x*_*ij*_ denotes the edge community assignment of the edge between nodes *i* and *j*. The *i*th column of **X**, which we denote as **x**_*i*_ = [*x*_1*i*_, …, *x*_*Ni*_], encodes the community labels of all edges in which node *i* participates. Note that we do not consider self-edges, so the element *x*_*ii*_ is left empty.

From this matrix, we can compare the edge communities of nodes *i* and *j* by calculating the similarity of vectors **x**_*i*_ and **x**_*j*_. Here, we measure that similarity as the fraction of elements in both vectors with the same community label. That is:

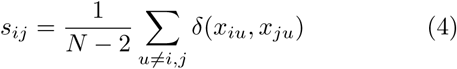

Here, *δ*(*x, y*) is the Kronecker delta, and takes on a value of 1 when *x* and *y* have the same value, but is zero otherwise. Note that the scaling factor is *N* − 2 because we ignore the self-connections *x*_*ii*_ and *x*_*jj*_. Repeating this comparison for all pairs of nodes generates the similarity matrix, **S** = {*s*_*ij*_}.

### Modularity maximization

In the main text, we computed system and whole-brain edge community similarity matrices. To discover the meso-scale structure of these matrices we used a multi-scale modularity maximization algorithm [23, 40, 41]. Modularity maximization detects meso-scale structure according to a simple principle: clusters are groups of nodes whose actual connection weight is greater than what we would expect by chance. This general framework is flexible and, through parameterization can be used to detect clusters of different sizes [39] and across layers (time [91], subjects [92], frequencies [93]).

Formally, the modularity quality function is expressed as:

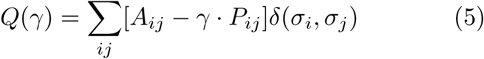

where *A*_*ij*_ is the observed weight of connections between nodes *i* and *j, P*_*ij*_ is the expected weight under some null model, *γ* is a structural resolution parameter, and *δ*(*x, y*) is the Kronecker delta and is equal to 1 when the community assignments of nodes *i* and *j*, denoted as *σ*_*i*_ and *σ*_*j*_, respectively, are identical and is equal to 0 otherwise. The inclusion of the delta function means that the double summation is over node pairs that fall within communities. Thus, *Q*(*γ*) measures the total weight of within-community connections less their expected values. The modularity maximization framework seeks to maximize the value of *Q*(*γ*) by selecting nodes’ community assignments.

Here we used a uniform null model, i.e. *P*_*ij*_ = 1 for all node pairs. Combined with the resolution parameter, *γ*, communities detected under this null model represent groups of nodes whose average similarity of edge community profiles exceeds *γ*. Note that we selected this particular null model deliberately, as previous studies have shown that it is especially well-suited for networks whose weights reflect statistical measures of similarity or correlation [40, 41]. We further note that this null model has been used in previous studies [33, 94–97].

In more detail, we selected 200 values of *γ*, linearly-spaced over the interval [0, 1]. At each value, we ran a Louvain-like algorithm to optimize modularity [98, 99]. Because this optimization algorithm is non-deterministic, we performed 50 iterations at each value of *γ*. We then aggregated *γ* values into 10 linearly-spaced intervals and, within each interval, used to detected clusters to generate a single representative set of clusters using a consensus clustering algorithm [42]. Briefly, this algorithm involved estimating the co-assignment matrix from the detected clusters, whose elements indicate the fraction of times that nodes *i* and *j* were assigned to the same cluster across all partitions within that interval. We then calculated the expected fraction (by randomly permuting nodes’ community assignments independently for each partition). The observed and expected co-assignment values can be used to define a consensus modularity function that we optimized using the same Louvain-like algorithm (1000 repetitions). If *any* of the 1000 partitions were dissimilar, we recomputed a co-assignment matrix and the expected co-assignment and repeated the algorithm. These two steps – calculation of co-assignment values and clustering – were repeated until convergence, i.e. all detected partitions are identical. In practice, the algorithm converged in three or fewer iterations.

### Edge community segregation

In the main text, we described a procedure in which we imposed edge community structure onto eFC matrices and measured a quantity that we referred to as an index of “segregation”. To calculate the segregation index, we measured two quantities induced by edge communities: *eFC*_*within*_ and *eFC*_*between*_, which measure the average eFC weight within and between edge communities. The segregation index, then, is simply the difference in these two quantities:

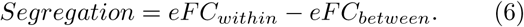

Because we define edge communities to be groups of edges with similar co-fluctuation patterns, we expect *eFC*_*within*_

### Differential identifiability

Suppose we had a dataset comprising many scans from many subjects. We would say that subjects are “identifiable” if, given a scan’s worth of data from one subject, we could accurately identify other scans from the same subject [44]. This intuition can be formalized using the measure *differential identifiability* [43]:

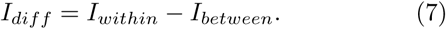

In this expression *I*_*within*_ and *I*_*between*_ are the mean similarities among scans from the same and different subjects. Here, we measure similarity as the Pearson correlation between regions’ edge community similarity vectors. Thus, *I*_*diff*_ measures how much more similar subjects are to themselves then they are to other subjects.

---

**FIG. S1.**
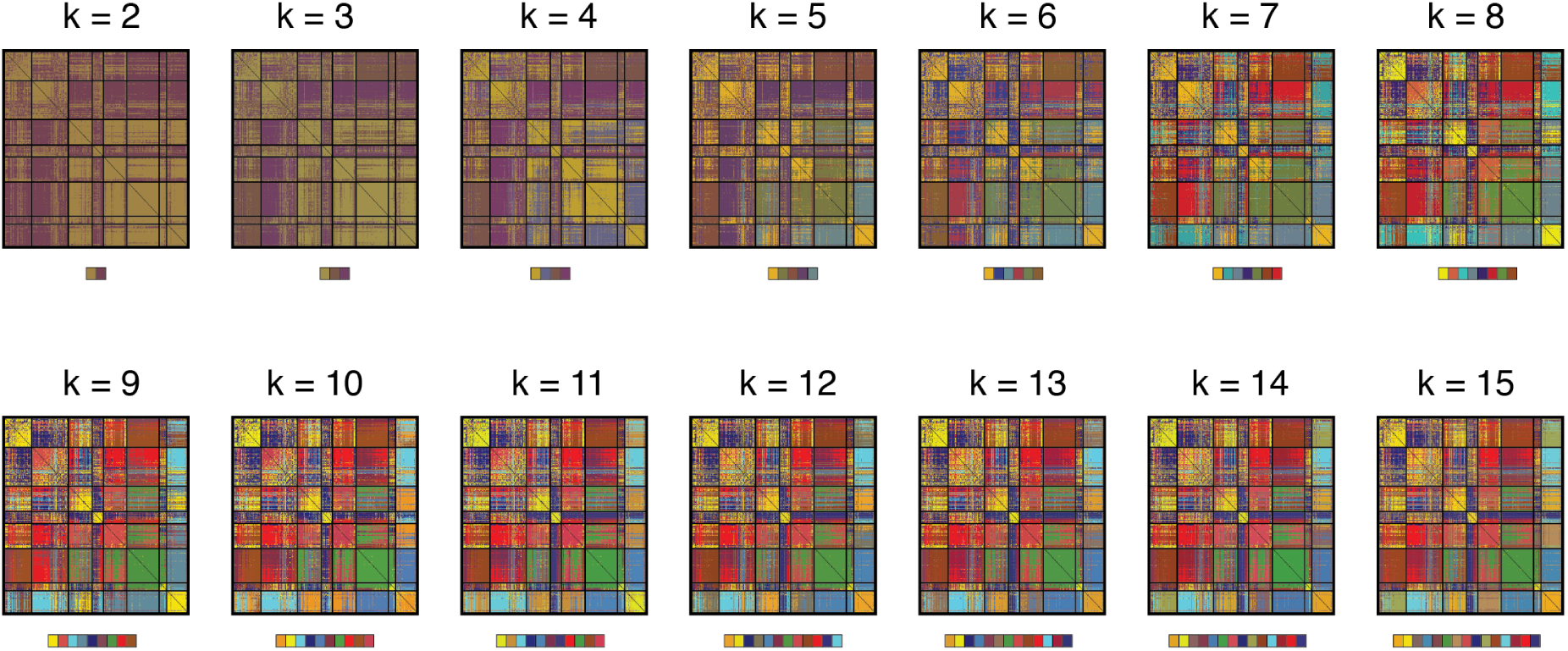
Edge communities mapped into node-by-node matrices.

**FIG. S2.**
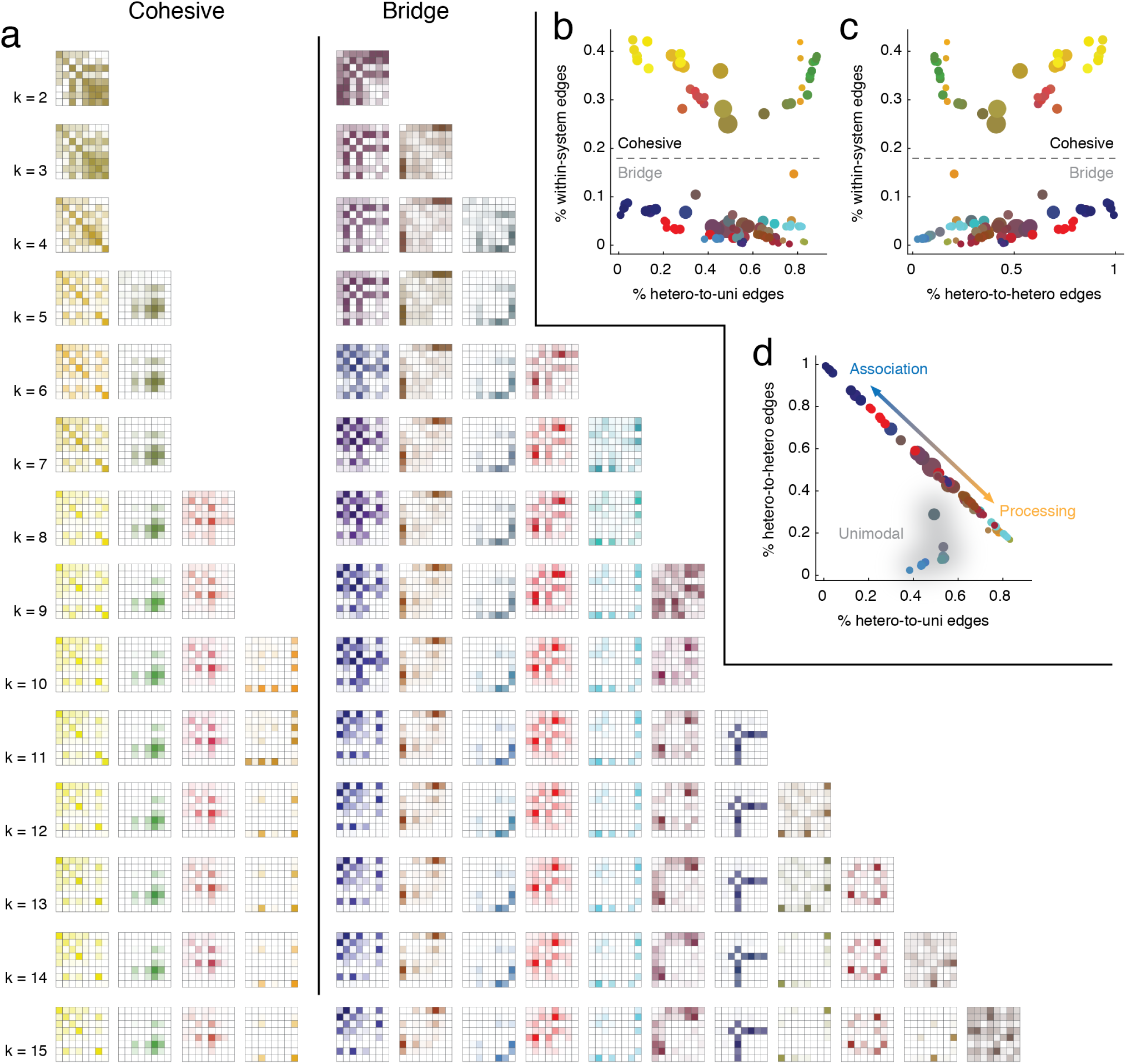
Edge community templates and classification with different numbers of edge communities. In the main text we described a procedure for classifying edge community “templates” – system-level representations of edge communities – as “cohesive” or “bridge” communities and further sub-classifying bridge communities as “processing” or “association” bridges. The main text included results with the number of edge communities fixed at *k* = 10. Here, we show results with different numbers of edge communities, ranging from *k* = 2 to *k* = 15. (*a*) Edge community templates at different *k*. In panels *b, c*, and *d*, we compare different features of edge templates and find evidence at all *k* of cohesive, bridge/processing, and bridge/association communities. In panels *b* and *c*, we plot the fraction of edges associated with each edge community that fall within the same system (y-axis in both plots) *versus* the fraction of edges linking heteromodal systems to themselves and to unimodal systems. These plots emphasize the division between cohesive and bridge communities. In panel *d*, we focus only on between-system edges and show that bridge communities fall along a spectrum with pure processing and association communities at either extreme. We also find evidence of a small number of unimodal-to-unimodal edge communities made up of edges that link visual and somatomotor systems to one another.

**FIG. S3.**
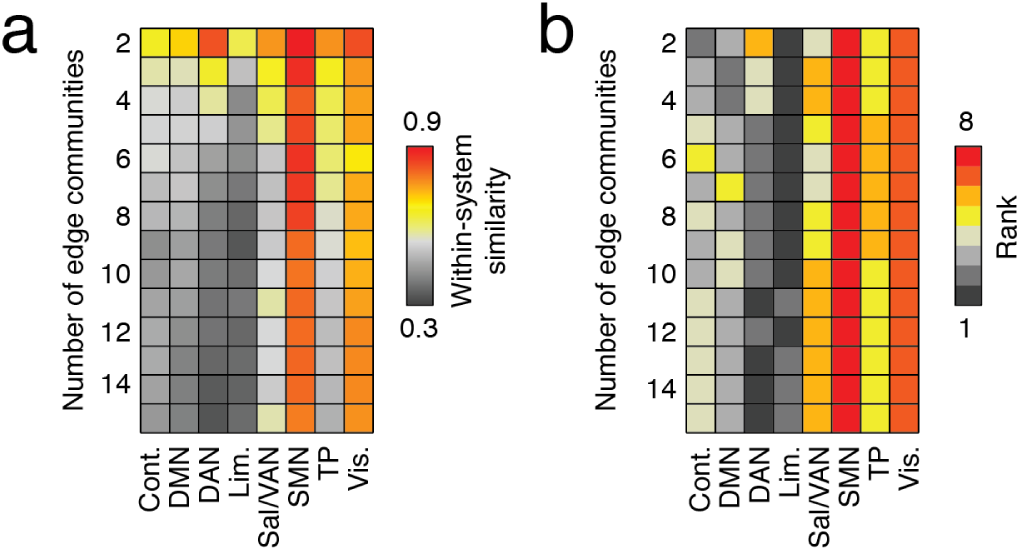
Mean within-in system similarity of edge communities profiles. In the main text, we showed that the within-system similarity of edge communities was greater in sensorimotor systems than in higher order systems. Those results were generated using a partition of the brain into *k* = 10 edge communities. Here, we show that this general pattern persists over a wide range of *k*. (*a*) Mean within-system similarity. (*b*) Rank-transform of mean within-system similarity.

**FIG. S4.**
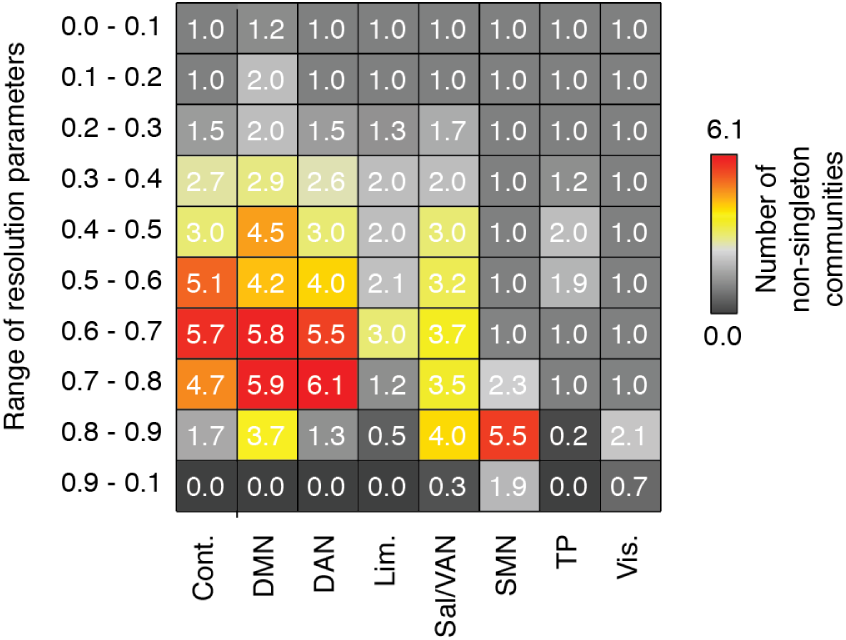
Number of non-singleton communities arranged by system. In the main text we clustered brain regions in every community based on the similarity of their edge community profiles to one another. The clustering algorithm was multi-scale, resulting in different cluster resolutions (from coarse clusters in which all nodes were assigned to a single cluster, to finer clusters where each node was assigned to its own singleton cluster). In the main text, we reported results at a single resolution. Here, we show mean results over all ten resolutions. With the exception of extremely fine partitions, these results are consistent with those presented in the main text. Namely, that higher-order cognitive systems are comprised of more sub-clusters than primary sensory systems.

**FIG. S5.**
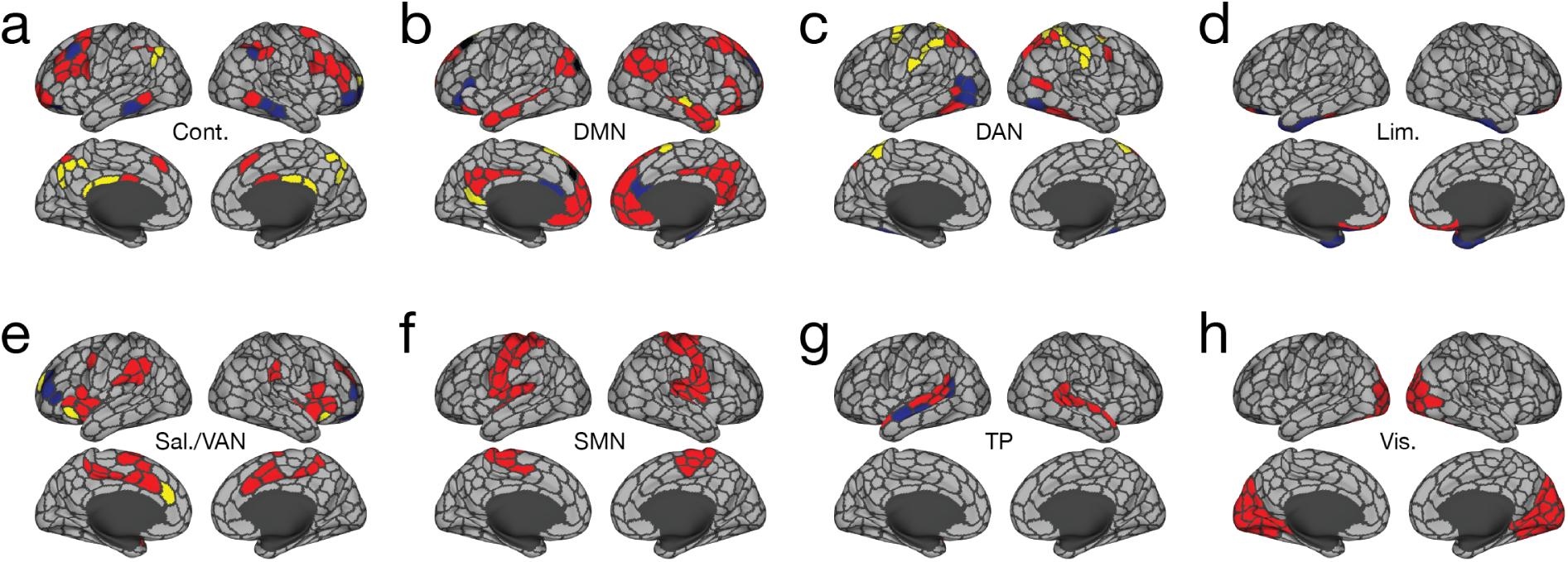
Sub-clusters derived from edge community profiles. We clustered regions belonging to eight brain systems based on the similarity of their edge community profiles. In the main text we investigated sub-clusters of the control network in more detail. Here, we show sub-clusters for all systems with the number of edge communities fixed at *k* = 10.

**FIG. S6.**
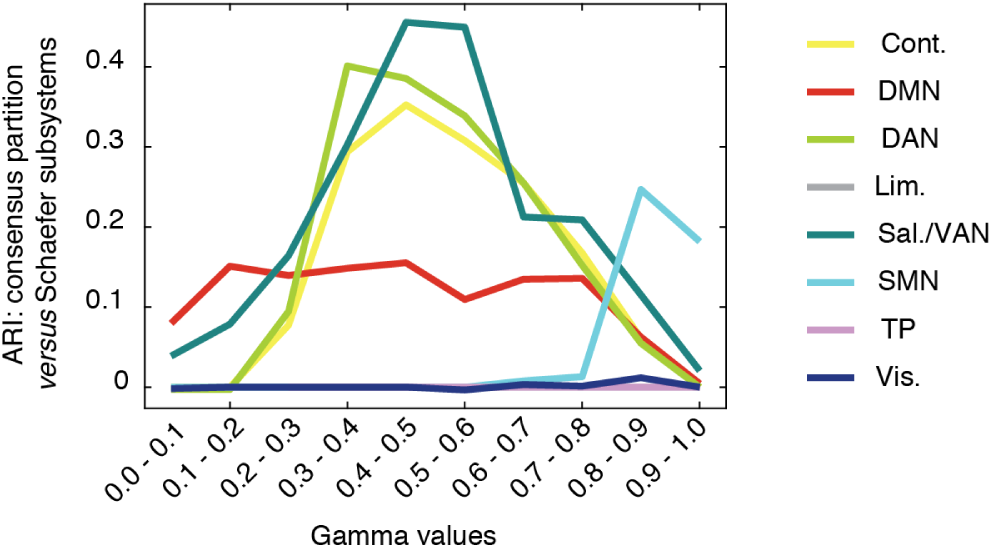
Comparison of detected sub-clusters with those from Schaefer/Yeo atlas. We compared the multi-scale clusters reported in the text with partitions reported in [38]. We used the adjusted Rand index (ARI) as a measure of similarity. Two identical partitions have an ARI equal to 1; two maximally dissimilar partitions have an ARI equal to 0. Here, we find that ARI for any system and at any scale never exceeds a value of 0.46 (Salience, Ventral Attention when 0.4 < *γ* < 0.5).

**FIG. S7.**
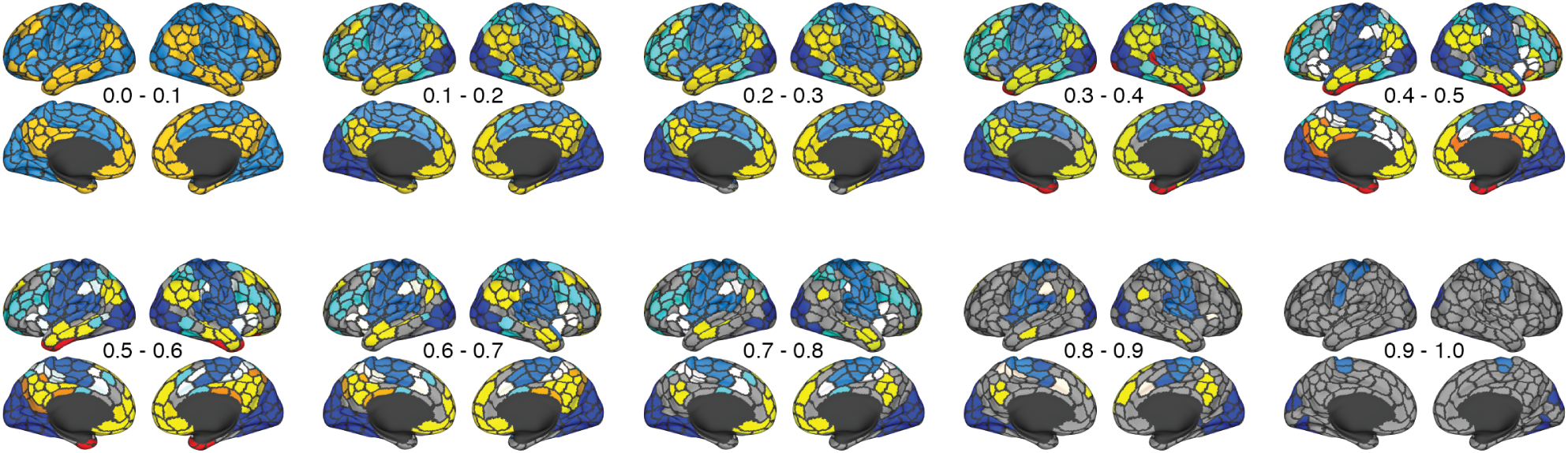
Whole-brain partitions generated using edge community similarity matrices with *k* = 10. In the main text, we described a partition of cerebral cortex into clusters based on the similarity of brain regions’ edge community profiles. We reported clusters detected at a single resolution (0.4 < *γ* < 0.5). Here, we show the full range of resolution. As in the main text, we aggregated small communities under a single label (gray).

**FIG. S8.**
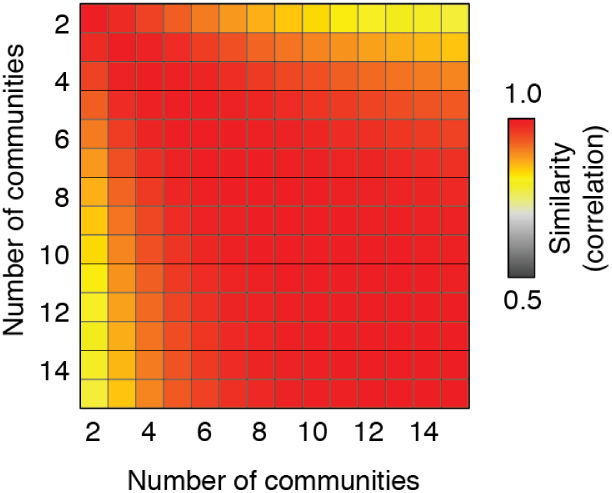
Similarity of Identifiability patterns across with different numbers of communities. In the main text we showed that when the number of edge communities was fixed at *k* = 10, regional identifiability was peaked in association cortex but lower in sensorimotor systems. We repeated that analysis with the number of communities ranging from *k* = 2 to *k* = 15, resulting in an identifiability score for each brain region. We then compared these patterns of identifiability across different numbers of communities and found that, on average, the similarity (Pearson correlation) was high: *r* = 0.96 ± 0.05. This observation suggests that over this range of *k*, the regions contributing to identifiability are highly similar.

